# The *α*C helix is a central regulator of PKR activation

**DOI:** 10.1101/2024.04.30.591909

**Authors:** Aaron G. Feinstein, James L. Cole, Eric R. May

## Abstract

Protein kinase R (PKR) functions in the eukaryotic innate immune system as a first-line defense against viral infections. PKR binds viral dsRNA, leading to autophosphorylation and activation. In its active state, PKR can phosphorylate its primary substrate, eIF2*α*, which blocks initiation of translation in the infected cell. It has been established that PKR activation occurs when the kinase domain dimerizes in a back-to-back configuration. However, the mechanism by which dimerization leads to enzymatic activation is not fully understood. Here, we investigate the structural mechanistic basis and energy landscape for PKR activation, with a focus on the *α*C helix – a kinase activation and signal integration hub – using all-atom equilibrium and enhanced sampling molecular dynamics simulations. By employing window-exchange umbrella sampling, we compute free energy profiles of activation which show that back-to-back dimerization stabilizes a catalytically competent conformation of PKR. Key hydrophobic residues in the homodimer interface contribute to stabilization of the *α*C helix in an active conformation and the position of its glutamate residue. Using linear mutual information analysis, we analyze allosteric communication connecting the protomers’ N-lobes and the *α*C helix dimer interface with the *α*C helix.

## Introduction

Human protein kinase R (PKR) belongs to the eIF2*α* family of protein kinases that inhibit translational initiation in response to different stress stimuli. Alongside a range of other roles in metabolic and apoptotic control, PKR functions centrally in the cellular innate immune response to infection by viruses^1–3^. PKR is activated by viral dsRNA and contains an N-terminal dsRNA binding domain and a C-terminal catalytic kinase domain (KD). Structural and biophysical analyses underscore a pivotal role for dimerization in PKR activation.^4^ The catalytic domain has the typical bilobal architecture found in protein kinases, consisting of a smaller N-terminal (N-) lobe and a larger C-terminal (C-) lobe.^5^ Upon binding viral dsRNA, the PKR kinase domains dimerize via their N-terminal lobe regions with the active sites facing away from each other in a back-to-back (BTB) geometry (**Fig. 1B**)^4,6^ This dimer configuration is believed to induce a prone-to-autophosphorylate (PTA) conformation^3,6^. PKR is then activated by autophosphorylation, and subsequently phosphorylates the translational initiation factor eIF2*α*, thereby blocking viral protein synthesis in the cell.

**Figure 1.**
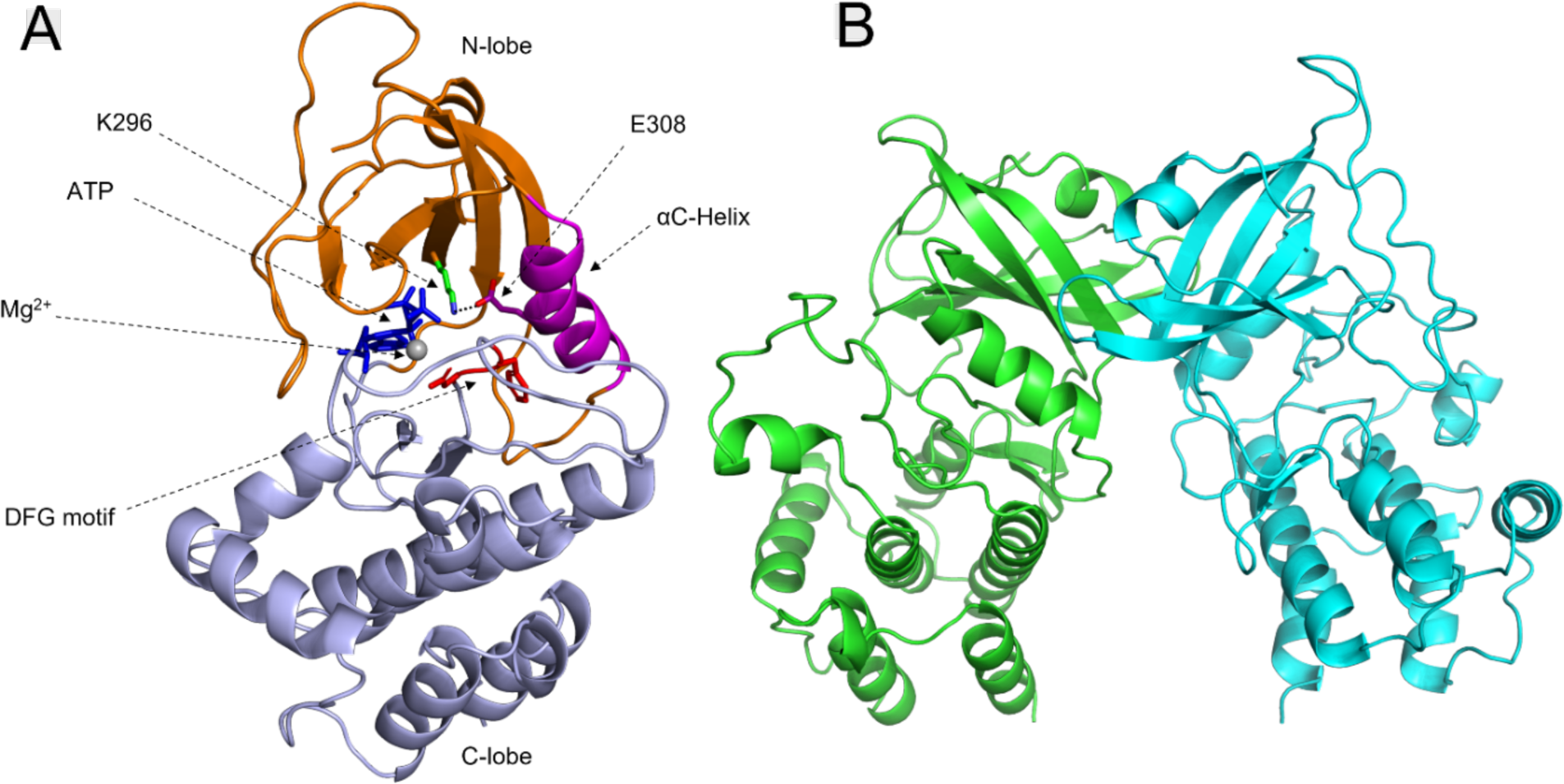
Structure of the PKR kinase domain. **(A)** The active site lies between the N-lobe (orange) and C-lobe (ice blue). The *α*C helix is shown in purple. In the active state, E308 of *α*CH forms a salt bridge with K296 which coordinates the *α* and *β* phosphates of ATP. D432 of the DFG motif coordinates a Mg2+ ion. **(B)** Structure of the back-to-back PKR kinase dimer, where the protomers are colored green and cyan (PDB ID: 63KL).

The PKR kinase domain comprises the structural elements that are crucial for enzymatic activity (**Fig. 1A**). The ATP-binding site is positioned between the N- and C-lobes. In the C-lobe, an aspartate located with the DFG motif coordinates a magnesium ion. In many kinases the DFG motif adopts alternative active (DFG-in) and inactive (DFG-out) conformations. In DFG-in conformations, the DFG aspartate’s sidechain faces inward; i.e., the sidechain is pointed toward the catalytic center of the KD, where it coordinates one or more Mg^2+^ atoms which in turn orient ATP for phosphoryl transfer. Conversely, in DFG-out conformations, the aspartate sidechain faces away from the ATP binding site. Generally, the DFG motif has a ratchet-like action, when the aspartate sidechain faces in, the phenylalanine faces out and vice versa. Another conserved regulatory element of kinases is the *α*C helix (*α*CH). In the active state, E308 of *α*CH forms a salt-bridge with K296 on *β*3, which in turn coordinates the *α* and *β* phosphates of bound ATP. Displacement of *α*CH in active kinases can induce disruption of this salt-bridge. Spanning the N- and C-lobes is the hydrophobic regulatory spine (R-spine)^7^, consisting of residues L312, Y323, and F433 in PKR^8^. In active kinases, the sidechains of these residues tend to be stacked linearly, whereas they are unaligned and often separated in inactive kinases.

The formation of the BTB dimer is a key step in the activation of PKR^3^. Interfacial residues that form interdimer salt-bridge and hydrogen bonding interactions are highly conserved amongst eIF2*α* kinases and mutations that disrupt these interactions block PKR kinase activity^9,10^. However, the mechanism by which dimerization induces the adoption an active conformation remains undefined.

In this study, we employ molecular dynamics (MD) simulations of the PKR kinase domain to compute a free energy profile and analyze the allosteric mechanism for dimerization-induced activation. We show that the *α*CH conformation functions as a central structural modulator of activity in PKR, while the conformation of the DFG motif is not predictive of activation status. Using an enhanced sampling approach, we calculate the free energy profiles for activation for both monomeric and dimeric PKR. We demonstrate that the displacement of *α*CH away from the catalytic center (and the resultant inactivation) is more probable for monomeric than dimeric PKR. Additionally, we perform linear mutual information analysis (LMI)^11^, where we observe correlated inter- and intramolecular dynamics connecting the dimer interface to the rest of the kinase domain. Finally, we perform contact mapping analysis to structurally analyze the dimer interface and to determine how the BTB configuration can stabilize the *α*CH and promote catalytic activity.

## Methods

### Structure modeling and molecular dynamics simulations

We conducted MD simulations (**Table 1**) of the kinase domain of active-state PKR in dimer and monomer forms using an x-ray crystal structure as our initial state (PDBID: 6D3K, chains B and C). In this structure, two PKR kinase domain monomers form a BTB dimer which induces an active-like conformation in the absence of autophosphorylation. Both chains were used in BTB dimer simulations, while only the chain C was used for monomer simulations. Unresolved loop regions (chain B: 255, 334-355, 439-451; chain C:334-356, 441-444, 449) were modeled using Schrödinger Prime v.1^12,13^ For clarity, throughout the manuscript, we refer to the dimer as composed of chain A and chain B, where chain A refers to PDB B chain structure and chain B refers to the PDB C chain structure. The completed structures were energy minimized using the Desmond package^14^ with the OPLS3^15^ force field and reached convergence after 1,460 steps. For simulations of ATP containing complexes, the existing ADP was replaced by ATP using UCSF-Chimera, v.4^16^ and manually adjusted with Schrödinger Maestro v.5^17^. There are no extant inactive structures of the PKR kinase domain: existing structures either have a phosphorylated activation loop and are thus active or are unphosphorylated but still in an active-like conformation. Unphosphorylated monomeric PKR is generally considered to be inactive, and we generated a homology model (**Fig. S1**), based on an inactive crystal structure of a closely related eIF2α kinase, GCN2 Homology models of the inactive PKR monomer were generated with I-TASSER^18–20^, using the inactive GCN2 kinase structure (PDBID: 1ZYC), as a template. Compared to the active-state monomer used here, the modeled inactive PKR monomer KD includes 3 additional N-terminal residues (YTV--) and 12 additional C-terminal residues (--KKSPEKNERHTC).

**Table 1.** Simulation times and descriptions.

| Sim | No. Windows | Simulation Time | Total Simulation Time | Simulation Type | # Atoms | Box dimensions (nm) | PDBID; Chain | Quaternary Structure |
| --- | --- | --- | --- | --- | --- | --- | --- | --- |
| 1 | NA | 927 ns | 927 ns | Equilibrium | 71,567 | 8.87 X 8.87 X<br>8.87 | 6D3K; C | Monomer |
| 2 | NA | 973 ns | 973 ns | Equilibrium | 65,027 | 8.59 X 8.59 X<br>8.59 | Homology Model based on template PDBID: 1ZYC | Monomer (inactive homology model) |
| 3, 4 | 24 | 129, 108 ns | 5.7 μs | WEUS | 118,266 | 10.50 X 10.50 X<br>10.50 | 6D3K; B,C | Dimer |
| 5, 6 | 24 | 114, 108 ns | 5.3 μs | WEUS | 65,955 | 8.64 X 8.64 X<br>8.64 | 6D3K; C | Monomer |

Total time, all simulations: 12.9 μs

In total, six simulations systems were analyzed for this study (**Table 1**). All-atom MD simulation inputs were generated using CHARMM-GUI^21,22^ and GROMACS 2019^23^. Throughout, simulations were solvated with TIP3P waters^24^ and ionized with 150 mM NaCl. GROMACS 2019 with the CHARMM-36m^22^ forcefield was used for all production simulations. In all cases, a cubic simulation box was used. Energy minimization of the dimeric and monomeric equilibrium systems used a steepest-descent algorithm. All systems underwent 100 ps of NVT equilibration followed by 100 ps of NPT equilibration, at 300 K and 1 atm. All production runs of the MD simulations were performed in the NPT ensemble, with the pressure isotropically maintained at 1 atm and the temperature at 300 K. Temperature coupling was controlled with a velocity rescaling stochastic thermostat^25^, with a 1.0 ps time constant. Pressure coupling used the Parinello-Rahman scheme^26^, with a time constant of 5.0 ps. All simulation steps utilized particle mesh Ewald summation for calculating long-range electrostatics, with a Fourier spacing of 1.2 Å. For Van der Waals and short-range electrostatics, GROMACS’ force-switch shifting function was applied at the cutoff distance of 10 Å to smoothly reduce the interaction potentials to zero at 12 Å. The production runs of the simulations were performed with a 2 fs timestep, with frames saved every 10 ps.

### Geometric analysis of PKR’s DFG motif and *α*CH

In the absence of a solved inactive structure of PKR, we sought to identify a regulatory structural element that was likely to modulate PKR’s activity. Toward this end, we focused on two of the kinase motifs most frequently considered predictive of catalytic activity: the DFG motif and *α*CH. We developed a program based on the KinaMetrix web-based software^27,28^. KinaMetrix uses a machine learning algorithm, trained on an extensive annotated set of kinase structures, to classify DFG and *α*CH conformations as active and inactive by scoring them with geometric descriptors, using an active structure of the archetypal kinase, protein kinase A (PKA), as a reference. These descriptors include an extended description of the DFG motif, as well as *α*CH angles and dihedrals. Our program was designed to continuously monitor the change in these metrics over the course of several μs long MD trajectories (based upon PKR active structures and inactive homology models), rather than scoring the descriptors of a single structure. We use a similar comparison algorithm to score DFG status, and a simplified version of the KinaMetrix *α*CH analysis which is based on the orientation of the vector pointing from the C-terminal end of the helix to its N-terminal end. We normalized this *α*CH axial vector to a superposition of the *α*CH on an active PKA structure. Any deviation in the angle or center-of-mass (COM) position of the helix produces a value < 1 (i.e., an angle to the PKA vector that is < 0°). The DFG scoring criteria are based upon the ratchet-like character of the conserved kinase DFG motif, whose flipping in and out generally involves coordinated rotation of the aspartate and the phenylalanine. In the *DFG-in* conformation, the DFG aspartate faces the ATP binding cleft, and the phenylalanine points away from it. This conformation is required for effective ATP docking and proper orientation of its phosphate moieties. The canonical *DFG-out* conformation is the reverse, in which the phenylalanine faces (in fact, blocks) the nucleotide binding pocket, and the aspartate faces away. Hence *DFG-in* is considered an activating conformation, while *DFG-out* is inhibitory.

Here, a query kinase’s DFG state is defined by two parameters, D1 and D2, describing vectors orthogonal to the alpha carbon (C*α*) to gamma carbon (Cγ) vector of the aspartate (D1) and phenylalanine (D2) side chains **(Fig. S2)**. Negative values denote a relative angle greater than 90°; positive, less than 90°. Thus, positive values of D1 indicate an aspartate vector more like that of PKA’s (i.e., facing the catalytic pocket, or DFG-in); positive values of D2 indicate a phenylalanine vector more like PKA’s (i.e., facing away from the catalytic pocket – again, DFG-in). The negative corollary indicates a DFG-out state. The query molecule’s DFG state is given by two comparison vectors for its aspartate 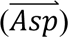 and phenylalanine 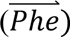 (**Eq. 1-2**).

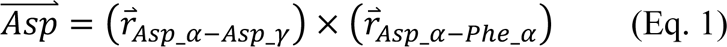

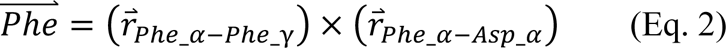

Where 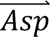 is defined as the cross product of the vector describing the query aspartate sidechain’s angle and the vector pointing from C*α* of the DFG aspartate to the phenylalanine C*α*. 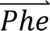 is similarly given by the cross product of the query Phe sidechain vector and the vector between the Asp and Phe C*α*. These sidechain-orthogonal vectors are used instead of vectors directly representative of sidechain orientation because the orthogonal vectors are insensitive to rotation of the sidechains in the same plane as the DFG backbone. DFG indicators D1 and D2 are then scored based on the normalized dot product of the query molecule’s vectors with those of the reference molecule, PKA (**Eq. 3-4**),

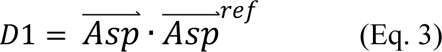

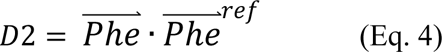

Where D1 can be taken to represent the relative angle between PKR and PKA’s aspartate sidechain, and D2 their phenylalanine sidechains. Four 1-μs scale equilibrium simulations were compared using this method. *α*CH orientation, D1, and D2 were compared in 2D scatterplots of the trajectories. Systems 1 and 2 (**Table 1**) were used to generate **Fig. 2-3**.

**Figure 2.**
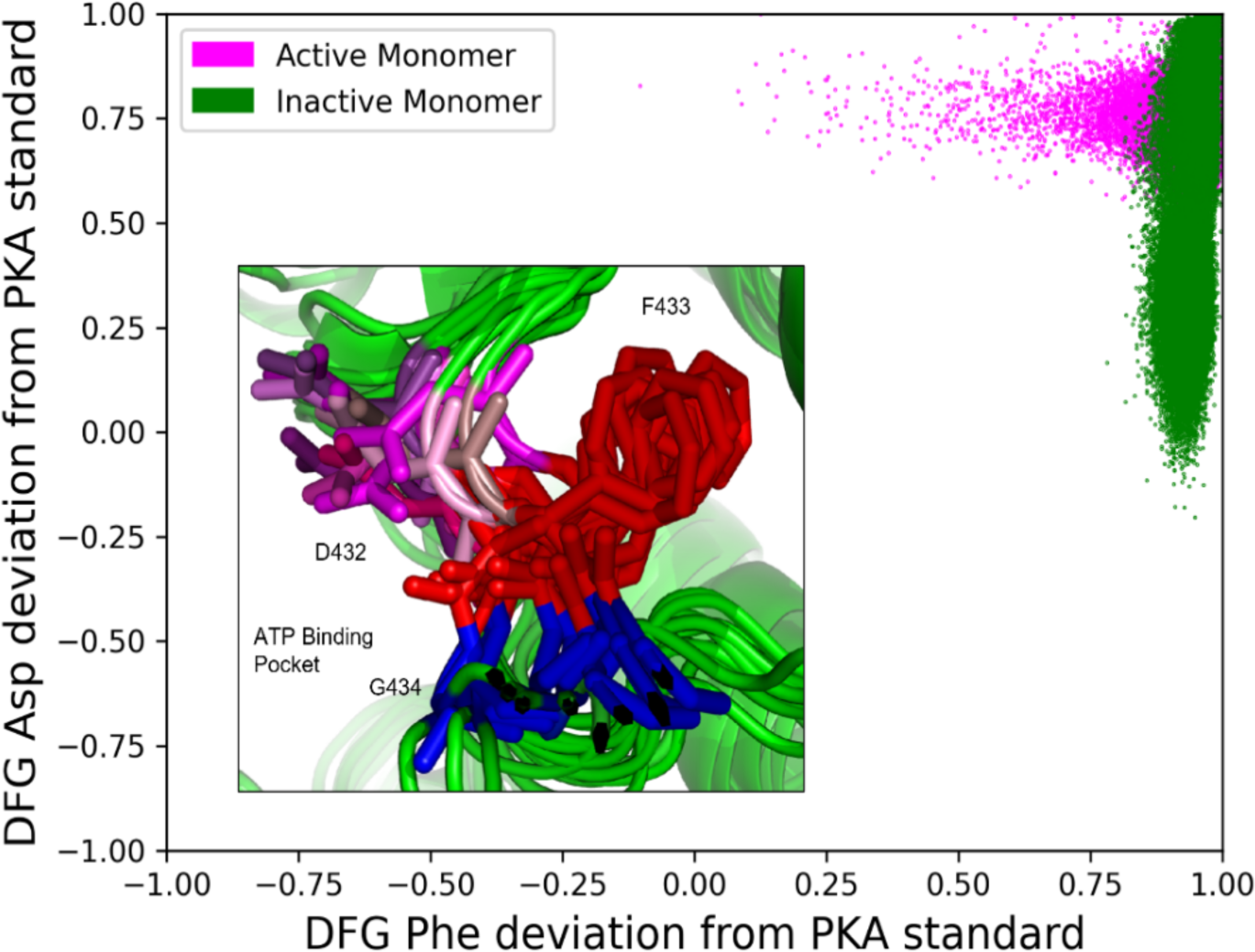
Geometric analysis of the DFG motif. Each data point represents a frame (every 10 ps) from an MD simulation. Indicators are plotted in magenta for the active kinase, and green for the inactive kinase. Negative values indicate conformations with the aspartate facing away from the catalytic center or the phenylalanine facing inward, corresponding to inactive states. **Inset:** Ten superimposed structures from an apo (no ATP or Mg ligands) ∼1 µs equilibrium MD simulation of inactive PKR kinase. D432 adopts a variety of conformations but does not undergo the characteristic flip corresponding to the canonical DFG-out state. F433 remains outside the nucleotide binding site.

**Figure 3.**
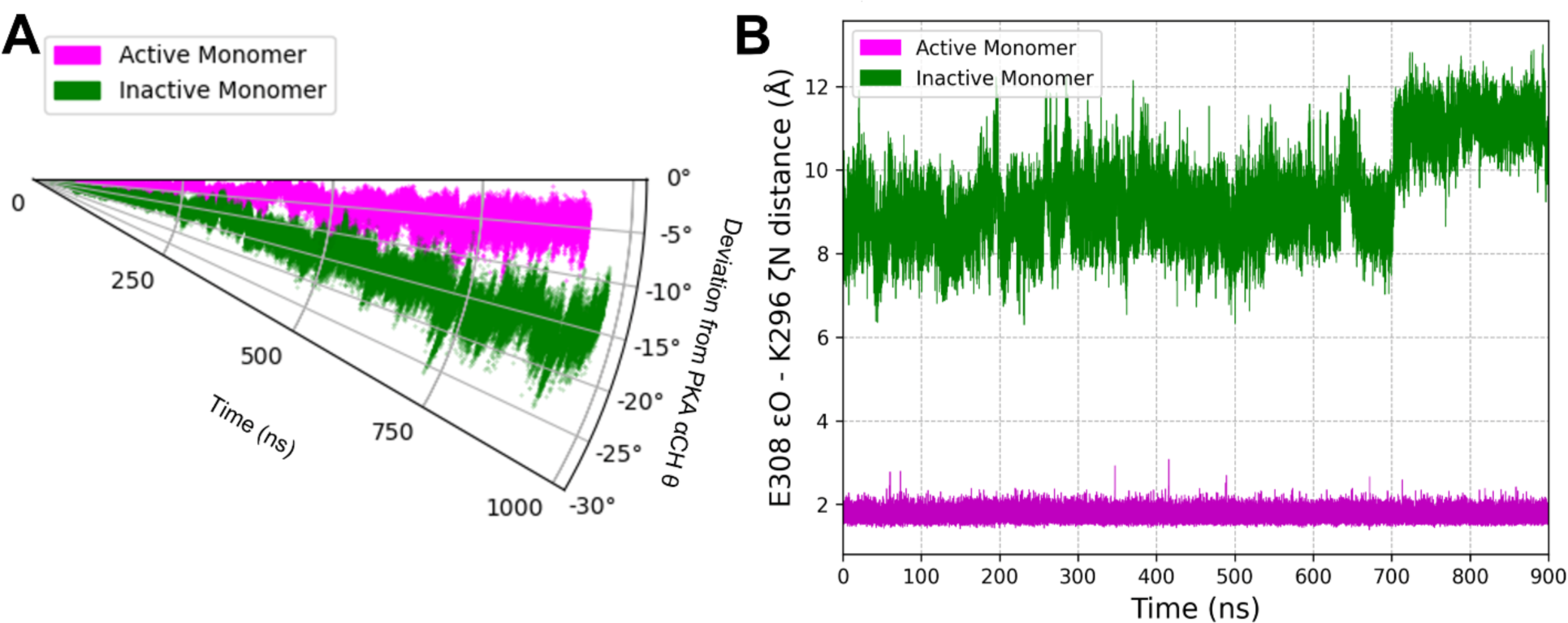
Geometric analysis of the conformation of the αC helix. **(A)** The deviation of the axial αCH vector from an active PKA reference in ∼1 µs simulations of active (magenta) and inactive (green) PKR kinase domain. Each point represents the angle from the reference in a single frame. **(B)** Distance between E308 and K296 in ∼1 µs simulations of active (magenta) and inactive (green) PKR kinase domain.

### Non-equilibrium molecular dynamics and window exchange umbrella sampling

We employed steered molecular dynamics (SMD) and umbrella sampling to compute free energy profiles of the displacement of *α*CH from and active(-like) conformation in both monomer and dimer states. In the dimer simulations, chain B *α*CH is displaced, which is the same chain used in the monomer simulations. Window exchange umbrella sampling (WEUS, i.e., Hamiltonian replica exchange) was performed to improve sampling of PKR’s conformational space along the reaction coordinate by swapping neighboring conformations according to the Metropolis criterion^29^. To drive the repositioning of the *α*C helix, SMD was performed with a potential applied between the atoms E308-εO (of *α*CH) and K296-ζN with a force constant of 2,000 kJ mol^−1^ nm^−2^, increasing the distance between the two residues from a starting point of 2.6 Å to approximately 14 Å at a rate of 0.05 Å/ns. This had the effect of displacing *α*CH away from the catalytic core, emulating a transition from an active to an inactive state. Simultaneously, a 1,000 kJ mol^−1^ nm^−2^ positional restraint was applied to the backbone atoms of ß-3, in the N-lobe, to prevent structural deformations during the SMD.

24 Frames along this reaction pathway were selected with 0.5 Å spacing, to serve as replicas (windows) for WEUS. In each window the E308-K296 distance was maintained with a harmonic distance restraint (force constant 2,092 kJ mol^−1^ nm^−2^). The 24 windows were run concurrently, with an exchange of replicas attempted every 1,000 steps (2 ps), based on a version of the Metropolis criterion:

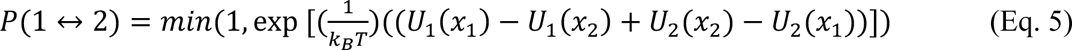

where *P* is the probability of exchanging two neighboring windows’ coordinates (*x_i_*), and *U* is the potential energy, including umbrella restraint, of a replica window. Window exchange probabilities ranged from 0.09 to 0.65, with an average exchange probability of 0.29 **(Table S1**). Potential of mean force (PMF) curves were computed using the GROMACS version of the weighted histogram analysis method (WHAM)^30^. Two independent SMD and WEUS simulations were run for each system (dimeric and monomeric) and the lowest-energy PMFs calculated for each system were compared. Error estimation was performed by block averaging of the WEUS data, where we computed four PMFs, each using 20 ns of data. PMF errors were computed by taking the standard error of the four 20 ns blocks for each system (**Figs. S3-S4**).

### Linear Mutual Information Analysis

We used the Bio3D R library^11^ to compute linear mutual information (LMI) matrices of the dynamics in cartesian space of backbone atoms in dimeric and monomeric equilibrium MD trajectories ∼1 us in length. LMI (unlike Pearson correlation methods, which measure linear relationships) considers both linearly related and orthogonal motions in its correlation calculations, returning a value of 0 (no correlation) to 1 (perfect pairwise correlation in time). LMI figures were plotted with the Matplotlib python library. Pairwise data from LMI matrices produced with Bio3D^31^ was used for the selection of putative allosteric pathways for further analysis. Contact mapping was performed using the GROMACS mdmat tool^32^.

## Results and Discussion

### DFG-In conformation is dominant in both active and inactive PKR monomers

Inactive or dysregulated protein kinases can adopt a wide array of conformations. This can make the identification of a central structural regulatory element (and identification of a collective variable (CV) for biased MD simulations) that reliably distinguishes between active and inactive conformations of the enzyme challenging. Several structural and dynamic features have been identified to characterize inactivation of kinases, including the disruption of the R-spine^33^, adoption of the DFG-out conformation of the DFG motif^34^, and displacement of *α*CH^35^.

Geometric analyses^27,28^ were performed on equilibrium MD simulations in which the orientations of PKR’s DFG aspartate (D432) and phenylalanine (F433) were compared to those of an active conformation of the archetypal kinase, PKA. In these analyses (**Fig. 2**), the dot product of the PKR DFG sidechain vectors with the PKA vectors was calculated; thus, (1,1) would indicate the DFG-in (active) conformation, whereas a value of (−1,−1) would indicate the DFG-out state (inactive). These analyses indicated low variability of PKR’s DFG motif, and both inactive and active PKR primarily exhibit DFG-in (active) conformations, though D432 demonstrates rare excursions into the negative regime. However, close examination of D432 dynamics shows that it is rarely, if ever, oriented such that its sidechain carboxyl would be unable to interact with Mg^2+^, which coordinates ATP. These non-canonical, intermediate DFG conformations would be unlikely to affect the activation state of the enzyme. F433 has even less mobility than D432. F433 is the main steric inhibitor of nucleotide binding in DFG-regulated kinases. As such, it is often called the “gatekeeper” residue^34^, due to its bulky sidechain ability to effectively block the docking of ATP to the catalytic site. While F433 remains in an out conformation, it is unlikely to have any effect on ATP binding. The lack of variability in DFG states between active and inactive conformations of the KD make this motif a poor predictive metric of PKR activation. Indeed, fewer kinases than previously believed reliably adopt the inactive, DFG-out conformation, calling the binary nature often ascribed to DFG conformation into question^27,34^.

### The *α*C helix is a highly dynamic structural feature of PKR

In contrast to the DFG motif, geometric analysis of the conformation of *α*CH showed a significant amount of variability in the inactive state simulation. By measuring the *α*CH axial vector’s deviation from a PKA reference we clearly observe its greater flexibility in our inactive model (**Fig. 3A**). In equilibrium simulations of the inactive PKR model, the orientation of *α*CH is quite dynamic. The active state αCH starts with an approximately 5° offset from PKA’s αC and doesn’t deviate more than 5° from that orientation. The inactive helix, however deviates from its initial orientation (already ∼15° off of the PKA standard’s axis) by as much as 10°.

The *α*CH also displayed significant bending, and translation away from the catalytic core (**Fig. S5**). In the inactive monomer, this has the effect of disrupting the formation of the E308– K296 salt-bridge. In contrast, *α*CH remained very stable in the active monomer in which the salt bridge was intact. On the simulation timescale, no interconversion between active and inactive kinase states was observed. However, the interatomic distance between E308 εO to K296 ζN in the inactive simulations varied by 5-6 Å (**Fig. 3B**). The demonstrable dynamic flexibility of *α*CH and its ability to moderate PKR activity by controlling this salt bridge make a strong case for its role as an essential regulatory feature of the kinase^7,13,35–38^. The *α*CH motif is universally conserved in Ser/Thr kinases^35^, and there is strong evidence that it frequently serves as a signal integration hub in protein kinases^36,38^. Residues from *α*CH interact with multiple catalytically important kinase motifs^33^, including the R-spine and elements of the activation loop. In PKR, the E308-K296 salt bridge is sensitive to the conformation and orientation of *α*CH, as E308 itself is positioned centrally along the helix.^8,39^ Hence, we chose *α*CH as the primary focus of our investigation to understand structural determinants and energetics of PKR activation.

### Dimerization stabilizes the active conformation of *α*CH

To probe the energetic coupling of dimerization with formation of an active kinase conformation we employed steered MD simulations of monomeric and dimeric PKR to generate active-to-inactive transition pathways. In these simulations, a force was applied to *α*CH to cause it to translocate, disrupting the E308–K296 salt-bridge. WEUS was performed on the transition pathways to obtain the potential of mean force (PMF) for the active to inactive transitions of monomeric and dimeric PKR (**Fig. 4, S4**). Extension of the E303-K296 distance by ∼8 Å requires approximately 5 kcal/mol for the dimer but less than 3 kcal/mol for the monomer, indicating that dimerization significantly stabilizes the active kinase conformation. Moreover, there is a prominent local minimum at ∼8 Å in the monomer which is not present in the dimer, resulting in an energy barrier of ∼1 kcal/mol to reestablish the salt-bridge in the inactive to active transition. The presence of the barrier in the monomer state indicates that the inactive conformation may be more stabilized compared to the dimer state, where the energy landscape of activation is primarily downhill. We therefore conclude that stabilization of a conformation of *α*CH that facilitates formation of the critical E303-K296 salt-bridge represents a primary mechanism by which dimerization promotes activation of PKR.

**Figure 4.**
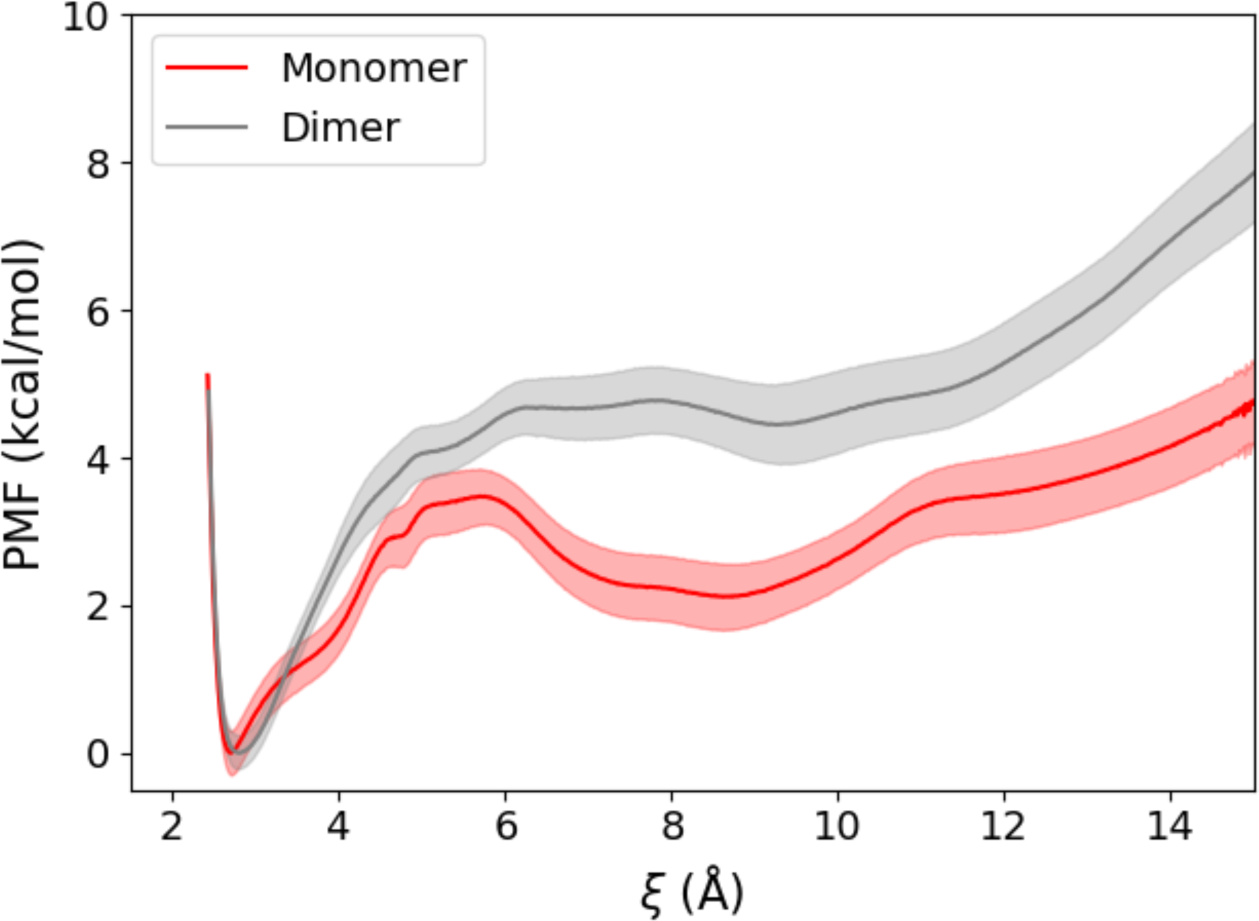
Potential of mean force (PMF) for αCH displacement. The dimer is in grey and the monomer is in red. The reaction coordinate (ξ) corresponds to the E308 εO - K296 ζN distance. The most favorable PMFs among the two trials for both monomer and dimer and shown. Shaded area represents the standard error of the mean computed by block averaging in 20 ns increments.

### Dimerization effects protein dynamics and correlated motions

Given that dimerization effects the energetics of αCH displacement (Fig. 4), which has strong implication on PKR activation, we next performed correlation analysis using linear mutual information (LMI) and contact mapping, to compare dimer and monomer states. **Figure 5A** shows the temporally correlated dynamics between residues within the PKR BTB dimer. This analysis captures both inter- and intra-protomer correlations, where a value of 0 corresponds to no temporal correlation, and 1 meaning the two residues are either fully correlated or fully anticorrelated in time. The upper left and lower right quadrants of the matrix represent inter-protomer correlations, while the upper right and lower left quadrants represent internal protomer correlations in chain B and chain A, respectively. Interestingly, LMI mapping shows an asymmetric pattern in the degree of intra-protomer correlation, with chain B (**Fig. 5A**, upper right quadrant) displaying considerably more coordinated dynamics than chain A (**Fig. 5A**, lower left quadrant). This may suggest that upon dimerization, the monomer that has the more rigid or coherent conformation is further stabilized. Alternatively, there may be an enzyme-substrate relationship between the protomers, in which a given monomer has adopted a conformation capable of preferentially stabilizing its partner upon binding, biasing that monomer toward autophosphorylation. Comparing the correlations in the B-chain of the dimer (**Fig. 5A**, upper right quadrant) to an inactive monomer simulation (**Fig. B-C**), we observe that the chain participating in the dimer has a distinct and generalized increase in intramolecular dynamic correlation of residues – the inactive monomer shows an average LMI of 0.44 while chain B of the dimer’s average LMI is 0.59. Possibly, dimerization has an overall coordinating effect on interfacing PKR protomers that accounts for this increase. Close residue packing at the dimer interface produces a greater degree of concerted dynamics, and this correlated motion propagates through the protomer. This is further evident when one chain is removed from its dimeric context (**Fig. S6A**). In terms of LMI, an isolated, active-like chain B shows a marked lack of intramolecular correlation after dissociation from chain A, emphasizing the role of dimerization in the synchronization of PKR’s dynamics. Presumably, given enough time under equilibrium conditions, an isolated active-like chain B would relax into a conformation conducive to more coordinated motion; the LMI of an inactive dimer under similar conditions (**Fig. S6B**) displays considerably higher correlations than active-like chain B in isolation. This is likely reflective of the fact that the conformation of the inactive monomer (in fact, any monomeric PKR KD that has remained isolated for a sufficiently long time), has equilibrated to an energetically favorable, if enzymatically inert, state.

**Figure 5.**
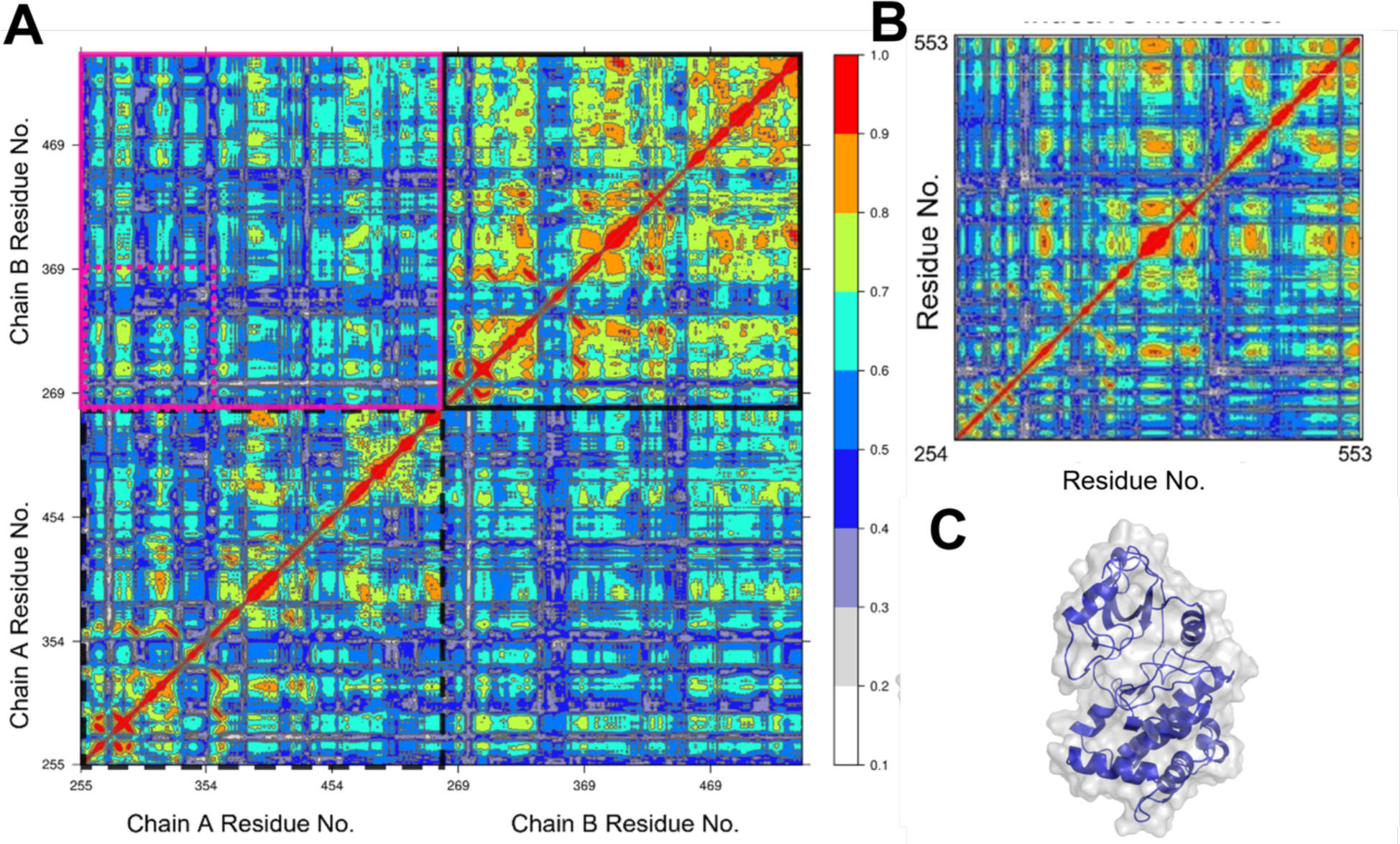
Linear mutual information analysis of PKR kinase. **(A)** LMI matrix of the PKR dimer. The dashed black boxed region represents dimer chain A, and the solid boxed region, dimer chain B. The colors indicate the degree of correlation, ranging from white (no correlation), to red (maximal correlation). The magenta box shows correlations between chains A and B. The small dotted magenta box demarcates correlations between the N-lobes of chains A and B. **(B)** LMI matrix of an inactive PKR monomer **(C)** Structure of the inactive monomer.

Examining the LMI of the N-lobe region of the dimer in more detail (**Fig. 6A**), four strips, or groups, of residues exhibiting strong interprotomer correlation (LMI values between ∼0.6 and ∼0.8) are apparent. Notably, these groups include several of the beta strands that comprise the majority of the N-lobe’s more rigid structural features: Group 1 contains ß1, Group 2 contains ß4, Group 3 contains ß2 and ß3, and Group 4 contains the N-terminus of ß5. Group 4, which averages <0.5 LMI, shows less interprotomer correlation than groups 1-3. This group, however, consists mainly of disordered loop residues so the lack of mutual correlation is perhaps unsurprising.

**Figure 6.**
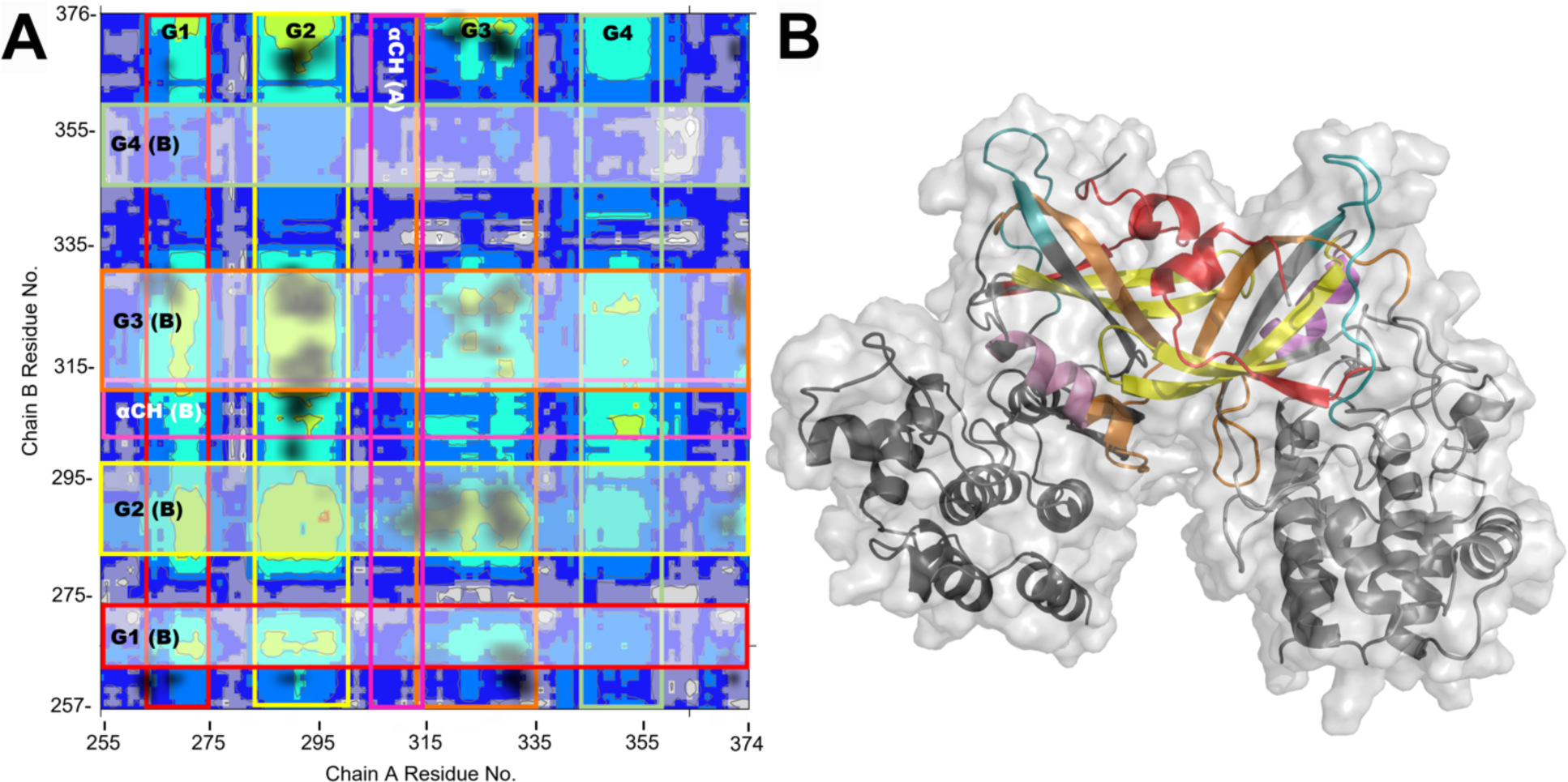
Inter-protomer dynamic correlation and contact map between PKR kinase domain N-lobes. **(A)** A close-up view of the dimer N-lobes inter-protomer correlations (dotted magenta box in Fig. 5A). Groups of residues in chain A and B that exhibit high correlation are shaded (horizontally, chain A) and/or outlined (vertically, chain B) in red, yellow, orange, and teal. αCH residues are outlined in magenta. Shadowed regions show areas of frequent direct contact. **(B)** Dimer structure colored by correlation group. Chain A is shown on the left, B on the right. Colored structural regions correspond to the groups of high interprotomer dynamic correlation shown in (A). Red: Group 1, residues T258-S275; Yellow: Group 2, residues F282-Y300; Orange: Group 3, residues L312-E335; Teal: Group 4, residues D345-K360; Magenta: αCH.

Residues comprising or adjacent to the interface between N-lobes have a high degree of mutual correlation, as well. Frequent contacts between interfacial residues with their conjugate protomer can be seen on the correlation map in the Group 2 and Group 3 regions, particularly where the A and B chains’ ß3 (R287-K299) and ß4 (N324-Y332) strands intersect, so high dynamic correlation at these sites is to be expected. ß1 (E271-G276), ß3, and ß4 all exhibit high interprotomer mutual information as well. In chain A, these beta strands also show strong correlation with chain B’s *α*CH. Chain A’s ß3, notably, shows the strongest correlation with chain B’s *α*CH. This is consistent with the fact that several residues on the amphipathic C-terminal end of the helix interact directly with hydrophobic chain A ß3 and ß2-ß3 turn residues: I288 and Y293. Indeed, it is possible that the close packing between hydrophobic residues of *α*CH and interfacial residues on the conjugate protomer is contributing to stabilizing the active conformation of *α*CH in the dimer. Although only a small number of residues make up this region of the interface, the hydrophobic effect could account for the ∼2 kcal mol^−1^ difference observed in the free energy profiles of the monomer and dimer^40^.

Of particular interest are the areas of the N-lobe LMI map (**Fig. 6A**) that exhibit high correlation in the absence of direct contact. While it might be unsurprising that cognate structures that exhibit frequent explicit residue contacts would be highly dynamically correlated, this correlation appears to propagate from C-terminal *α*CH residues at the interface (shown in orange in **Fig. 6A-B**) through ß4 and into ß2 and ß3 of both protomers, where no direct contact can be seen. Also notable, the long (Y332 through K358), disordered kinase insert Domain (KID) of chain A displays relatively strong correlation with chain B’s *α*CH as well. While no clear mechanistic role for the KID has been described thus far, there is some evidence that its deletion attenuates or abrogates kinase activity in PKR^41^.

Interprotomer correlation between cognate C-lobes (visible in **Fig. 5A**, as the upper right quadrant of the magenta boxed region) is high. In this region, residues rarely touch, but *α*F and *α*G (the binding site for PKR’s primary substrate, eIF2*α*) of chain A correlate strongly with several chain B N-lobe structures: its *α*O helix (residues Y261 through D266), ß3, ß4, and ß5. C-lobe to C-lobe correlations exist as well: the *α*F helices of both protomers share > 0.7 LMI; *α*F, *α*H, and *α*J of chain B, all show strong correlation with chain A’s *α*F helix. Slow, coordinated “clamshell” or “breathing” motions, in which the N- and C-lobes close around the KD active site, are frequently cited as a hallmark of enzymatic activity in kinases^42^. The high correlation between C-lobes may indicate that this motion is being captured by our analysis of the PKR dimer.

These pairwise atomic correlations point to highly synchronized residue dynamics within and between the dimer protomers. The correlations may also suggest a dynamic communication pathway, coordinated by the N-lobes’ beta strands, which connects the protomers’ interface to more distal regions of the KD, such as the C-lobe helices. Concerted, large scale relative motions of the protomers may drive the coherent transmission of residue displacements linearly along the ß strands, maintaining or minimizing the distance between interfacing residues, and stabilizing the protomers’ active, more rigid conformation. Further, K296 is located in ß3, which, as mentioned, has high mutual interprotomer correlation. It is possible that, upon dimerization, the increase in outward-oriented correlated residue motion propagating from the interface and *α*CH through ß4 and into ß3 contributes to the stabilization of the critical E308-K296 salt bridge in chain B by synchronizing local dynamics. This effect appears to be asymmetric, with mean LMI above 0.6 for chain B’s *α*CH with chain A’s ß1 and ß3, but relatively low values for the reverse case – again, dimerization may preferentially affect the stabilization of one protomers’ *α*C helix or the other^35,36,38^.

## Conclusions

Geometric analyses of multiple microsecond scale MD simulations of active PKR and an inactive PKR model demonstrated that the kinase does not adopt the canonical DFG-out conformation, indicating that this element is not be a central regulator of activation (**Fig. 2**). In contrast, *α*CH shows higher levels of conformational flexibility in the inactive model than in the active state (**Fig. 3**). Along with this, its roles in phosphoryl transfer chemistry and R-spine assembly make it a compelling candidate for PKR’s regulatory hub.

Using enhanced sampling methods, we calculated the free energy of displacement of the *α*CH in a PKR dimer and monomer. The resultant PMFs revealed a greater free energy difference between active and inactive states for the dimer, demonstrating that dimerization stabilizes the active conformation of PKR (**Fig. 4**).

Correlation analyses suggest that stabilization and coordination of residues and motifs within the protomers of a PKR dimer may occur asynchronously after they interface, rather than simultaneously. Contact mapping points to hydrophobic interactions between *α*CH and the dimer interface as a possible explanation for the differential free energy for *α*CH displacement in the monomeric and dimeric configurations of PKR. These findings also suggest that several of the N-lobe beta strands may serve as allosteric conduits connecting the *α*C helix to distal residues, and that allosteric communication along this structure is markedly increased once PKR has dimerized (**Figs. 5-6**).

Taken together, our results establish an explicit mechanistic link between dimerization and activation, mediated by *α*CH. As a future experimental approach, the beta strands implicated in communication between the BTB dimer interface and *α*CH could be subjected to mutational scanning and accompanying activity assays. Future *in silico* experiments, such as displacing the chain A αCH in the dimer to explore the notion that one protomer acts a “substrate” could also be merited. Moreover, the suggestion of a dynamic link between the kinase insert domain and *α*CH might justify a closer examination. This unusually long loop domain could be more functionally relevant than currently realized and has been generally neglected in the literature. Here, we have demonstrated that *α*CH is a pivotal element in PKR activation. While our findings shed new light on kinase regulation, they also underscore that there is terrain still left to explore to fully understand these complex biological systems.

## Acknowledgements

This work was supported by the National Institutes of Health (Grant R35-GM119762 to E.R.M.) Computational resources for this work have been provided through the University of Connecticut Storrs HPC center.

