## Supporting Information for "The *α*C helix is a central regulator of PKR activation"

Supporting information contains: supplemental figures S1-S6 and supplemental table S1.

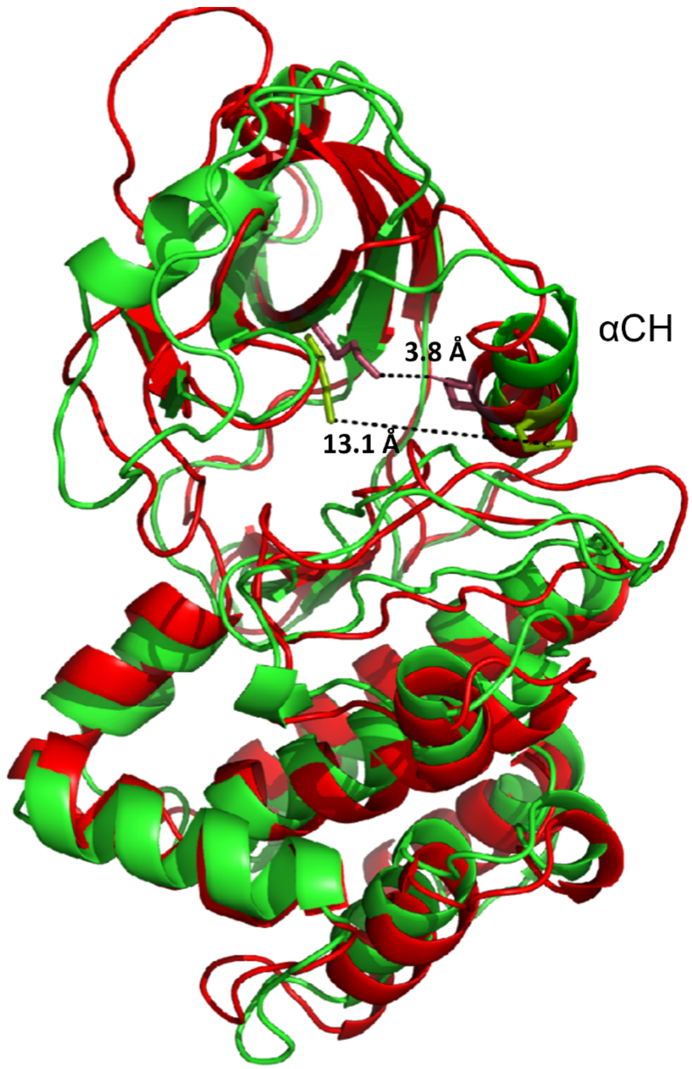


**Figure S1. Comparison of the structures of inactive and active PKR kinase.** The model of inactive PKR kinase (green) is superimposed on active PKR kinase (PDBID: 6D3K; red). The E308- K296 distances are indicated. Helix *α*CH in the inactive structure is in a conformation with its N-terminal end pointing outward, thus disrupting the E308- K296 salt-bridge.

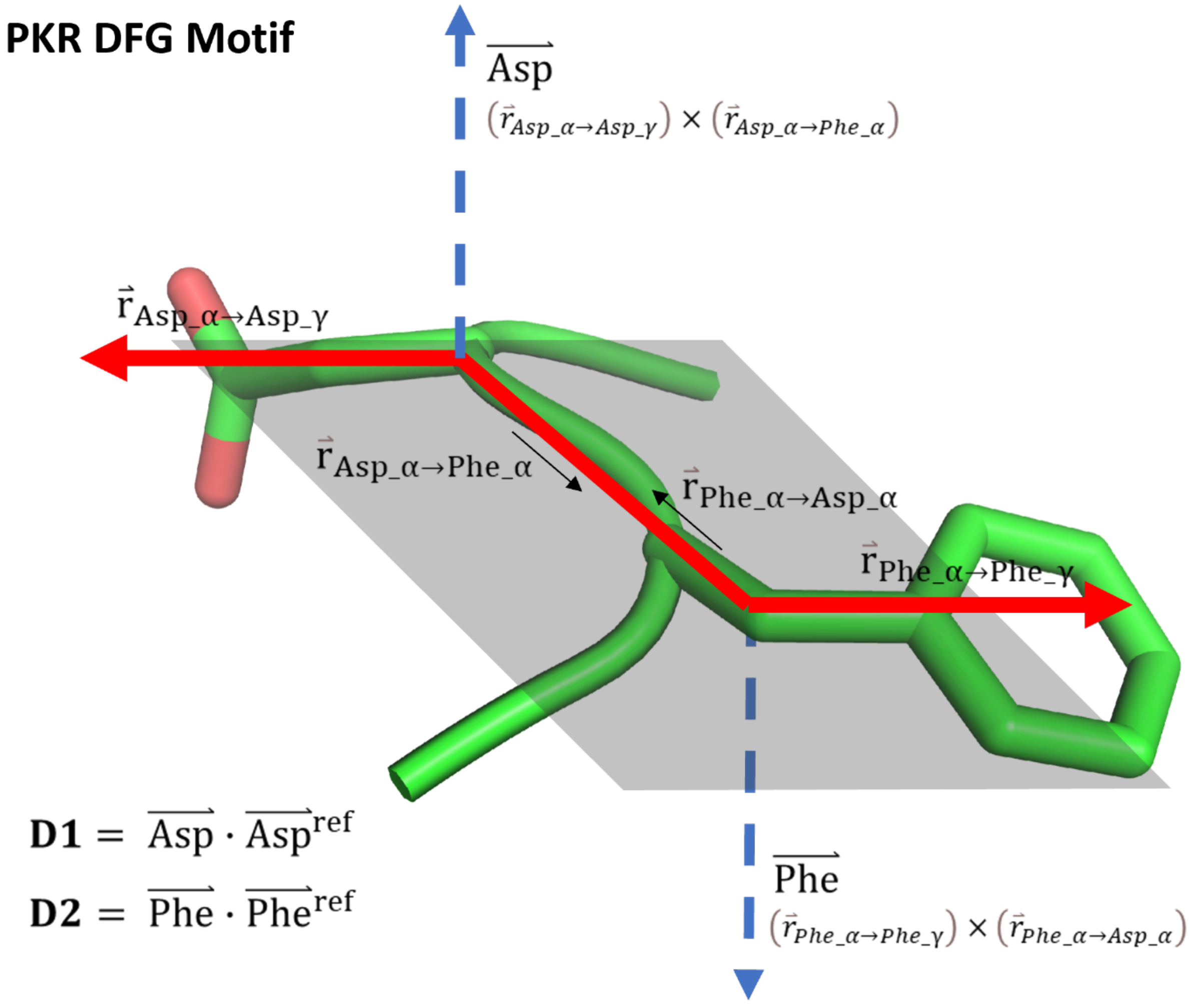


**Figure S2. Geometric Analysis of DFG.** PKR's DFG motif is scored with descriptors D1 and D2. The dot product between PKR and a PKA reference is calculated between vectors orthogonal to the plane described by the enzymes' Asp and Phe sidechains and the bond between them.

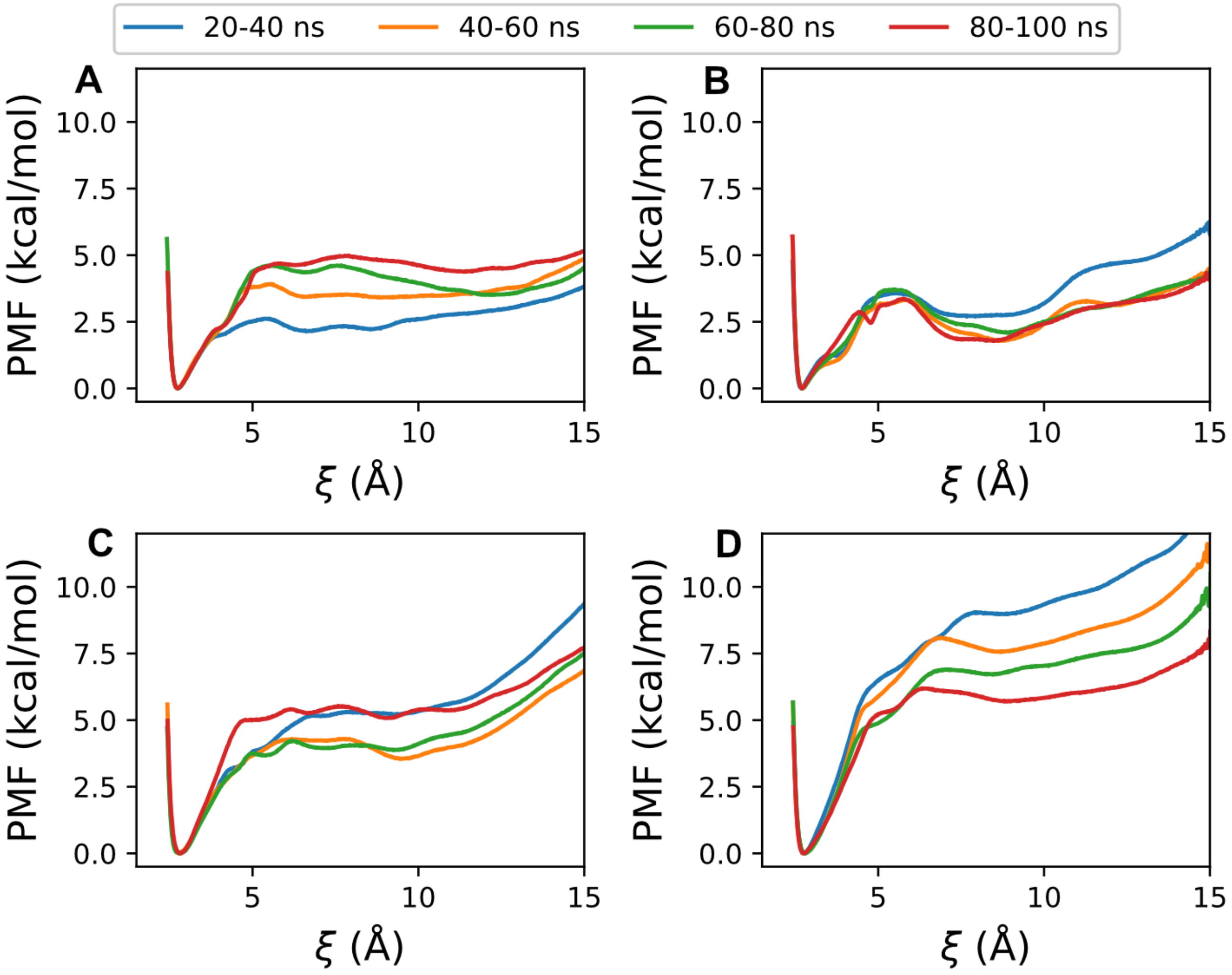


**Figure S3. Convergence of PMFs.** Block PMFs from WEUS are shown for monomer systems (**A-B**) and dimer systems (**C-D**).

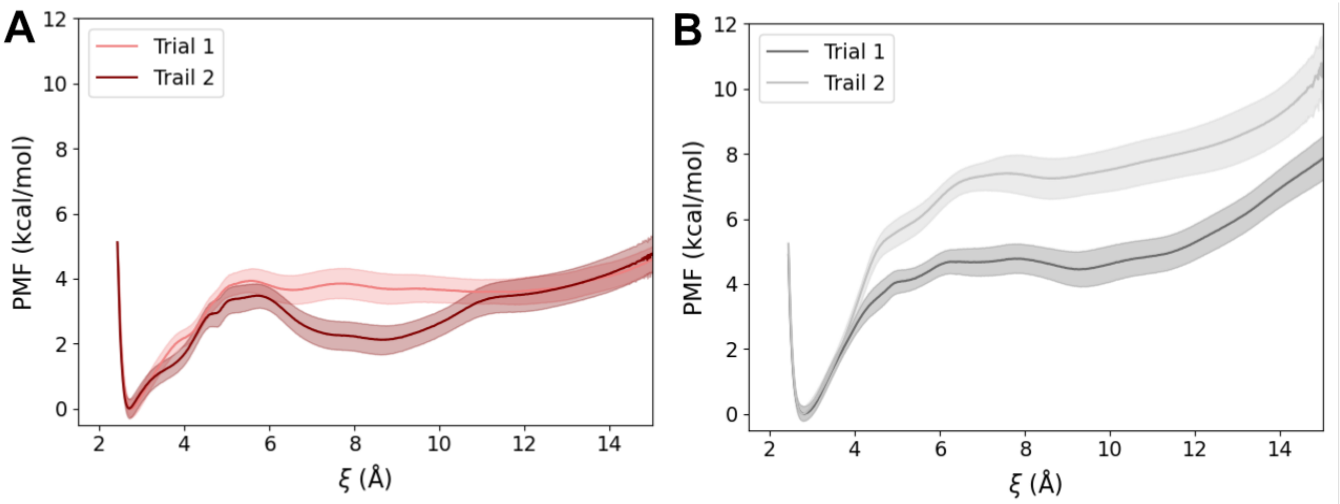


**Figure S4. WEUS replicate PMFs.** PMFs computed using WEUS from independent SMD trials for a monomeric system **(A)** and a dimeric system **(B)**. Error bars are the standard error computed over the four 20 ns blocks per system (block PMFs shown in **Fig. S3)**.

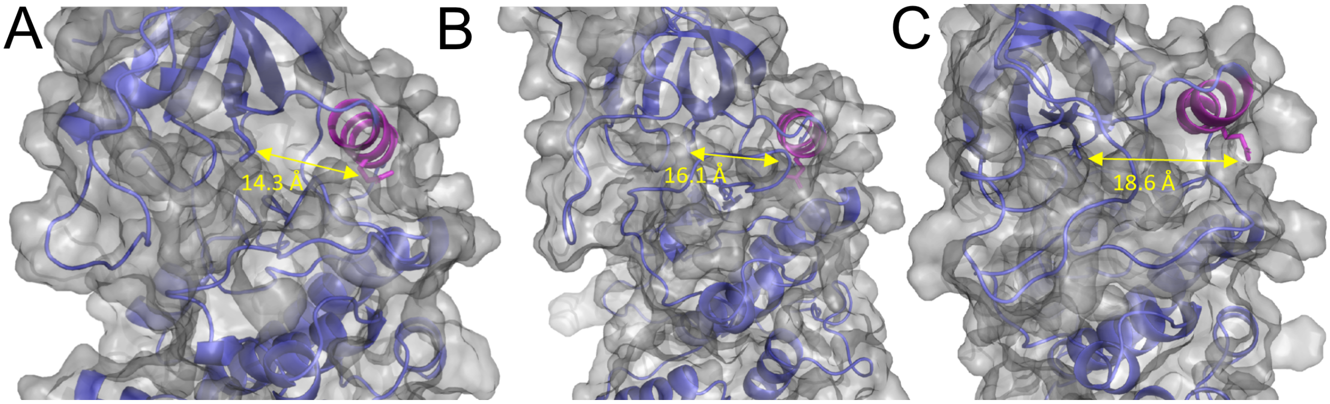


**Figure S5. Conformational fluctuations of αCH in inactive PKR. (A-C)** Aligned frames from an MD trajectory of inactive PKR kinase domain with a disrupted E308-K296 salt-bridge. The position and orientation of αCH (shown in violet) fluctuate considerably, with large concomitant variations in the E308 εO - K296 ζN distance.

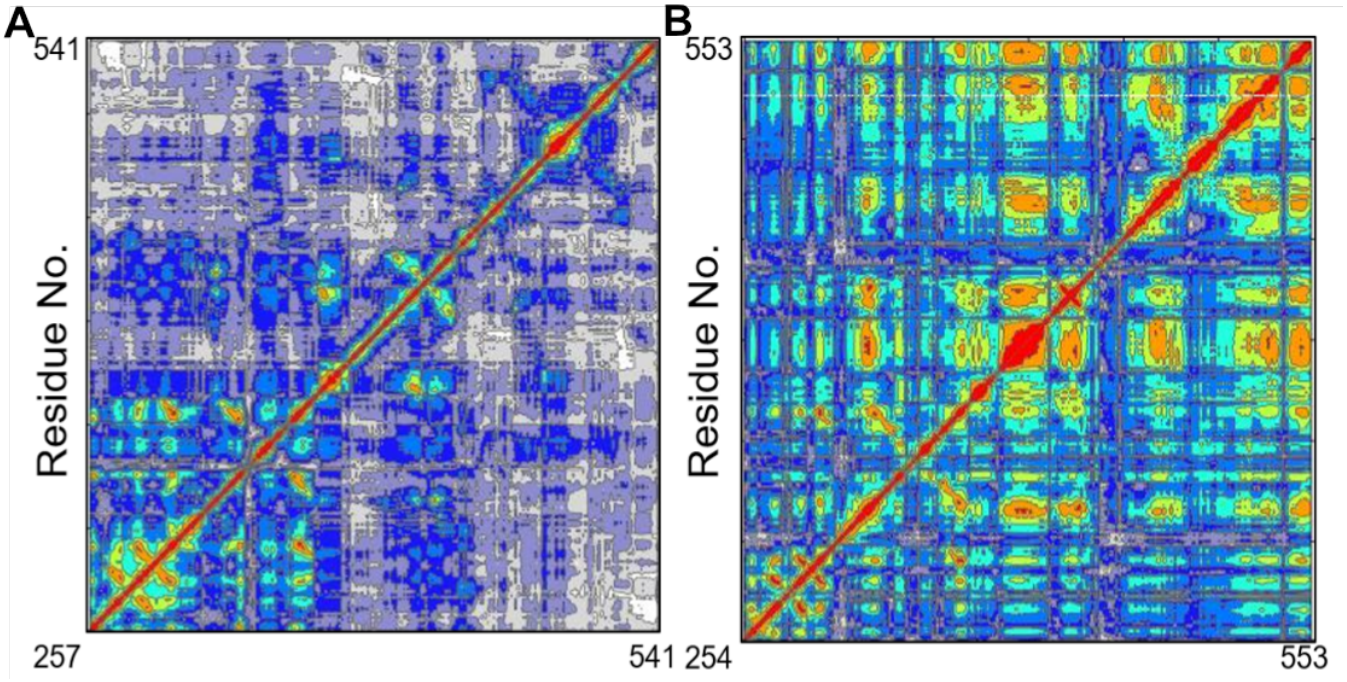


**Figure S6. LMI of isolated dimer chain B and inactive monomer. (A)** Separating the chain B protomer from its conjugate causes significant generalized decorrelation of its residues. **(B)** In the inactive model, residues are more correlated than the dissociated chain (**A**), suggesting that PKR KDs removed from their oligomeric state relax into more synchronous dynamics after a period of equilibration, though they do not attain the degree of correlation they display while participating in a homodimer.

**Table S1. WEUS replica exchange statistics.** Replica exchange probabilities for all HREX systems. Window exchange was done in GROMACS, while the PLUMED implementation of the Weighted Histogram Analysis Method was employed for PMF generation.

| System* | State | Run time (ns) | HREX WINDOW (Average probability of Hamiltonian exchange between neighboring umbrella windows) | | | | | | | | | | | | | | | | | | | | | | |
|---|---|---|---|---|---|---|---|---|---|---|---|---|---|---|---|---|---|---|---|---|---|---|---|---|---|
| | | | 0-1 | 1-2 | 2-3 | 3-4 | 4-5 | 5-6 | 6-7 | 7-8 | 8-9 | 9-10 | 10-11 | 11-12 | 12-13 | 13-14 | 14-15 | 15-16 | 16-17 | 17-18 | 18-19 | 19-20 | 20-21 | 21-22 | 22-23 |
| 5 | Dimer | 119 | 0.52 | 0.35 | 0.32 | 0.31 | 0.21 | 0.34 | 0.32 | 0.38 | 0.36 | 0.30 | 0.29 | 0.31 | 0.32 | 0.39 | 0.39 | 0.25 | 0.38 | 0.38 | 0.36 | 0.28 | 0.26 | 0.36 | 0.35 |
| 6 | Dimer** | 129 | 0.43 | 0.27 | 0.09 | 0.18 | 0.17 | 0.18 | 0.17 | 0.16 | 0.21 | 0.20 | 0.18 | 0.25 | 0.17 | 0.25 | 0.24 | 0.27 | 0.28 | 0.27 | 0.27 | 0.14 | 0.25 | 0.25 | 0.27 |
| 7 | Dimer | 108 | 0.63 | 0.50 | 0.46 | 0.42 | 0.19 | 0.26 | 0.34 | 0.27 | 0.30 | 0.35 | 0.36 | 0.38 | 0.32 | 0.30 | 0.38 | 0.39 | 0.36 | 0.37 | 0.35 | 0.36 | 0.36 | 0.38 | 0.39 |
| 8 | Mono | 187 | 0.60 | 0.49 | 0.24 | 0.38 | 0.37 | 0.21 | 0.37 | 0.64 | 0.57 | 0.64 | 0.60 | 0.61 | 0.61 | 0.30 | 0.32 | 0.30 | 0.30 | 0.18 | 0.44 | 0.31 | 0.31 | 0.16 | 0.54 |
| 9 | Mono** | 114 | 0.60 | 0.43 | 0.34 | 0.26 | 0.23 | 0.27 | 0.24 | 0.30 | 0.35 | 0.37 | 0.32 | 0.31 | 0.32 | 0.33 | 0.31 | 0.29 | 0.31 | 0.30 | 0.31 | 0.31 | 0.32 | 0.32 | 0.31 |
| 10 | Mono | 108 | 0.46 | 0.23 | 0.16 | 0.10 | 0.13 | 0.20 | 0.20 | 0.17 | 0.18 | 0.19 | 0.19 | 0.18 | 0.13 | 0.17 | 0.19 | 0.20 | 0.20 | 0.18 | 0.20 | 0.21 | 0.21 | 0.18 | 0.13 |

*For system details, refer to Table 1
**Shown in Figure 4
